# Rank-based integration identifies convergent disease mechanisms across omics

**DOI:** 10.64898/2026.09.25.754380

**Authors:** Zifeng Qiu, Duncan Palmer, Luke Jostins, Andrew Lewis, Katherine Bull, Jagdeep Nanchahal, Yang Luo

## Abstract

The rapid growth of omics studies offers new opportunities to uncover disease mechanisms, yet findings from individual studies or single omics may be influenced by cohort-, platform– and layer-specific variation. Integrating evidence across studies and omics layers can reveal robust biological signals that are reproducible across molecular layers, but differences in assay platforms, sample sizes and feature coverage complicate direct combination of summary statistics. Here we present the Omics Rank-Based Integration Tool (ORBIT), a direction-aware framework for prioritizing concordant signals across omics layers using only within-dataset feature ranks and effect directions. ORBIT tests directional rank consistency across omics layers against a closed-form variance-gamma null distribution while accounting for inter-dataset correlation and incomplete feature overlap. Simulations show that ORBIT maintains well-calibrated false-positive rates under inter-omics correlation, increasing numbers of omics layers and missingness, while detecting concordant signals with increasing power as the number of layers grows. We applied ORBIT to tubulointerstitial transcriptomic data from patients with chronic kidney disease in the C-PROBE cohort and proteomic data from the Kidney Precision Medicine Project, identifying a concordant injury programme marked by inflammatory and fibrotic remodeling together with impaired energy metabolism. We also used ORBIT to integrate ten transcriptomic, four proteomic and one translatomic datasets in dilated cardiomyopathy heart tissue, identifying extracellular matrix remodeling and mitochondrial energy failure as the dominant concordant biological programmes. In both diseases, ORBIT prioritized concordant genes and identified pathways that were not significant in any single omics layer alone. These results show that directional rank integration can be used for integrating heterogeneous omics summary data and prioritizing robust molecular mechanisms.

## Introduction

Emerging high-throughput technologies have enabled comprehensive profiling of molecular phenotypes across multiple layers, including the genome, transcriptome, proteome and epigenome, at both tissue and single-cell resolutions [1, 2]. Integrating these datasets offers an opportunity to distinguish molecular changes that are reproducibly supported across studies and biological layers from those specific to individual datasets. Such concordant signals can provide stronger evidence for shared disease mechanisms and reveal biological processes that are incompletely captured by any single omics layer [3]. However, exploiting these rapidly growing omics datasets requires integration methods that can accommodate substantial heterogeneity in how molecular features are measured and analyzed. Integrating heterogeneous omics datasets remains challenging because studies can differ substantially in assay technology, sample size, feature coverage and statistical analysis. Methods that integrate individual-level data preserve detailed inter-feature structure and distributional information across studies, but are vulnerable to residual batch effects and incompatible data structures across datasets [4]. Summary-level integration based on effect sizes and P values is therefore a more practical alternative when individual-level data are unavailable or measurements are not directly comparable. Classical meta-analytic approaches, such as fixed– or random-effects models on log-fold changes, assume consistent study designs and comparable effect-size scales. Yet these assumptions are seldom met when integrating omics modalities [5]. P value merging methods such as Fisher’s combined probability [6], Brown’s method [7], directional P-merging methods (DPM) [8] and Stouffer’s methods [9] offer a more flexible alternative because they do not require comparable effect-size scales. However, when applied across heterogeneous omics layers, these methods can be influenced by systematic differences in statistical power and assay sensitivity, allowing layers that tend to produce smaller P values to dominate the combined results [10]. Furthermore, different studies may apply different statistical methods making the direct combination of summary statistics challenging.

Rank-based aggregation provides an alternative strategy by replacing absolute statistical magnitude with relative evidence within each dataset. Such approaches have been applied to differential-expression meta-analysis and gene-list integration, identifying features that are consistently highly ranked across replicate experiments or heterogeneous datasets. By operating on ranks, methods such as RankProd [11] and robust rank aggregation [12] reduce dependence on measurement scale and facilitate integration across experimental platforms without requiring joint normalization. However, existing rank-based approaches were developed primarily for integrating replicate experiments, differential-expression studies or ranked gene lists rather than heterogeneous multi-omics summary statistics. Therefore, these methods do not specifically account for background dependence between contributing datasets, differing feature coverage across molecular layers, or directional cancellation of discordant cross-omics effects.

To address these needs, we developed Omics Rank-Based Integration Tool (ORBIT), a direction-aware rank-based framework that integrates summary-level omics data to identify features with concordant signals across layers. By transforming omics association statistics into signed ranks, ORBIT focuses on the relative ranking and direction of each feature rather than effect-size magnitude or P value significance, allowing direct comparison across heterogeneous datasets. In simulations, ORBIT maintains a well-calibrated false-positive rate under between-dataset correlation and incomplete feature coverage, and gains power as omics layers are added. Applying ORBIT to chronic kidney disease (CKD) and dilated cardiomyopathy (DCM), we recovered coherent biological programs, including inflammatory and fibrotic remodeling with impaired tubular energy metabolism in kidney tissue, and fibrotic extracellular matrix (ECM) remodeling with suppressed mitochondrial function and fatty acid metabolism in DCM.

## Results

### Overview of ORBIT

The fundamental idea behind ORBIT is to identify features, such as genes, that are consistently ranked towards the same extreme across omics layers. Unlike methods that combine the strength of association across datasets, ORBIT does not ask how strongly a feature is associated with a phenotype, such as diseases, in each layer. Instead, it asks whether the feature shows consistent directional ranking across layers. ORBIT rewards consistent rankings and penalizes discordant ones, while a feature that ranks highly in only one layer retains its within-layer rank evidence. ORBIT therefore tests the integrated directional rank of a feature rather than the magnitude of its association within individual layers, making its statistical output distinct from existing methods.

ORBIT operates on summary-level data, requiring only feature rankings and their directions of effect from each omics dataset. Rankings can be derived from either directional or non-directional statistics, including P values, effect sizes or t-statistics. When ties occur, we break them at random. To ensure the same sign has the same biological meaning across omics layers, the direction of each feature is aligned using both its feature-level direction, such as up– or downregulation (*s_i_*_j_ ∈ {−1, +1} for each feature i in dataset j) and the reference direction of the corresponding dataset, such as case versus control. The aligned direction is defined as the product of feature-level direction *s_i_*_j_ ∈ {−1, +1} for each feature i in dataset j, and dataset-level reference direction 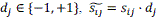.

We then transformed rank to place features on a common scale across datasets. For each dataset j, we normalized the rank, *r_i_*_j_, by the number of measured features, *m*_j_, giving *u_i_*_j_ = *r_i_*_j_/(*m*_j_ + 1), and transformed the normalized rank to a log-scale, *X_i_*_j_ = − log(*u_i_*_j_) (**Fig. 1A**). This transformation assigns larger values to features with more extreme ranks while making ranks comparable across datasets with different numbers of measured features. To accommodate incomplete feature coverage, we retained all features measured in at least one dataset and defined *A_i_* = 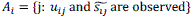, as the set of datasets in which feature i is observed. We then calculated the directional integration statistics, 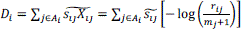 as the sum of signed log-ranks across observed layers.

**Fig. 1.**
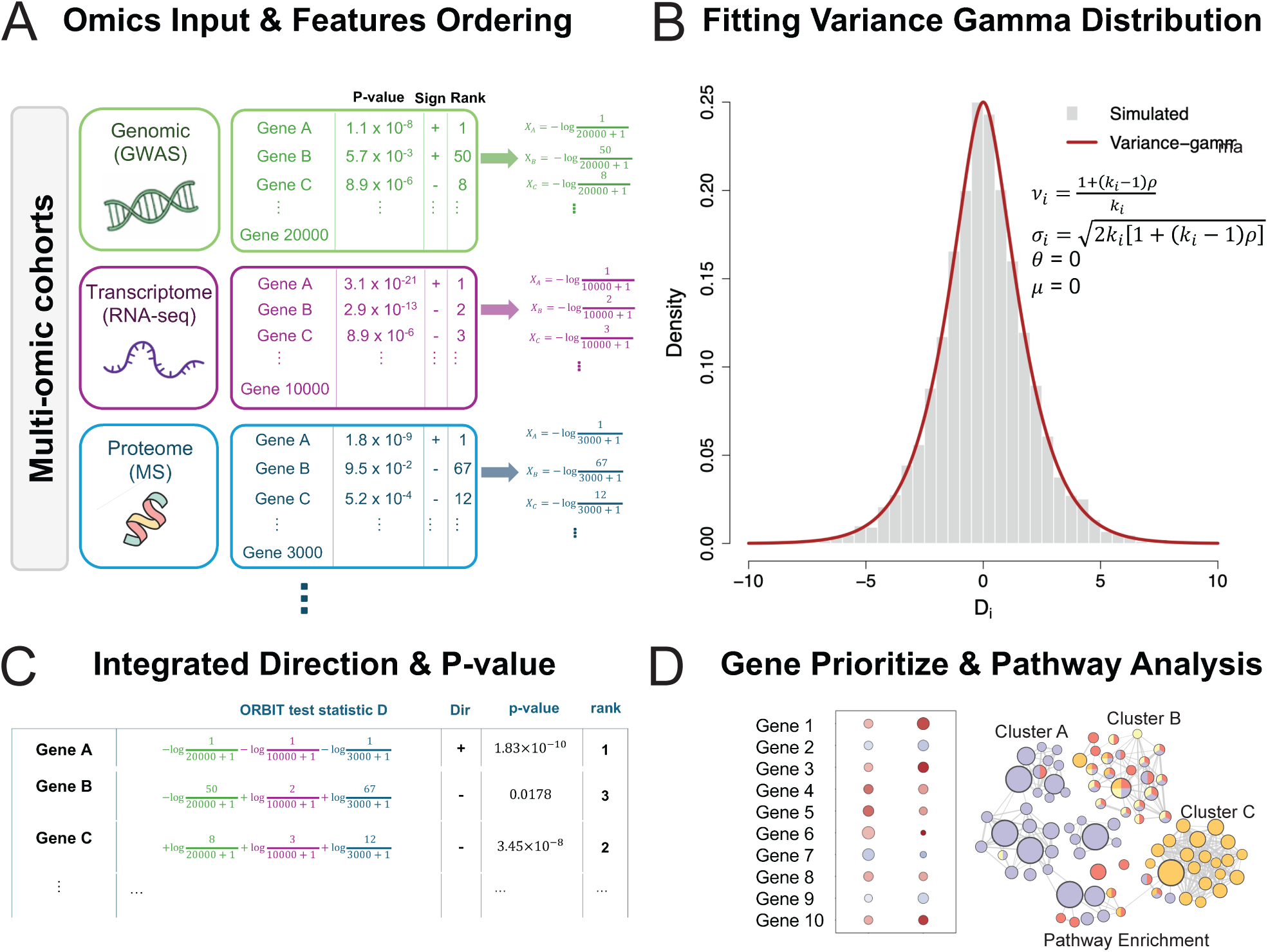
Overview of the ORBIT workflow. **(A)** Input from each omics layer, with feature directions and ranks derived from P values or other summary statistics, followed by log transformation of the ranks. **(B)** Fitting of the variance-gamma distribution and estimation of its parameters. **(C)** Calculation of the P value testing cross-omics directional rank and corresponding direction for each feature, followed by generation of the integrated ranking. **(D)** ORBIT output supports downstream gene prioritization and pathway enrichment analysis.

Under the null hypothesis, the integrated test statistic *D_i_* follows a symmetric variance-gamma (VG) distribution, which arises as the difference of two independent gamma variables. Following the parameterization of Madan et al. [13], the VG law is characterized by four parameters: a scale parameter σ that determines the dispersion, a shape parameter ν that determines tail heaviness, an asymmetry parameter θ, and a location parameter µ that shifts the center. Because the signed log-rank scores are symmetric at zero under the null, θ = 0 and µ = 0. Under independence, the remaining parameters are 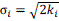 and 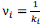, where *k_i_* = |*A_i_*| is the total number of datasets in which feature i is observed. To account for background correlation between datasets, we further adjusted σ and ν by moment matching using the estimated background correlation, ρ, among the datasets contributing to feature i, giving 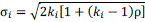 and 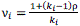 (**Fig. 1B**).

We ran simulations to validate the theoretical null distribution underlying ORBIT. We first evaluated the analytical variance-gamma distribution directly under the continuous-rank null. Monte Carlo simulations closely matched the analytical variance-gamma tail probabilities across the evaluated numbers of omics layers **(Supplemental Fig. 1)**. Because the derivation treats normalized ranks as continuous Uniform (0,1) variables whereas observed ranks lie on a finite discrete grid, we then compared the theoretical approximation with the exact or permutation-derived discrete-rank null. The continuous approximation was conservative, particularly with fewer ranked features, and converged toward the discrete null as the number of features and omics layers increased **(Supplemental Fig. 2)**, supporting its use as the ORBIT reference null.

ORBIT uses this theoretical null to assess the significance of each feature’s integrated directional rank. For each feature, the sign of *D_i_* indicates the direction of the concordant signal while the VG tail probability provides its statistical significance (**Fig. 1C**). The resulting integrated statistics and P values can then support both directional– and non-directional downstream pathway analysis, including Gene Set Enrichment Analysis (GSEA) [14] and over-representation analysis (**Fig. 1D**).

### Benchmarking ORBIT using simulated data

We evaluated ORBIT’s performance using simulated data. First, we considered four signal scenarios across two omics layers. In the first scenario, no feature carried signal (**Fig. 2A**). In the second, each omics layer contained independent, non-correlated signals (**Fig. 2B**). In the third, both layers contained correlated signals but in opposite directions (**Fig. 2C**). In the fourth, both layers carried concordant signals in the same direction, representing the signal ORBIT is designed to detect **(Fig. 2D)**.

**Fig. 2.**
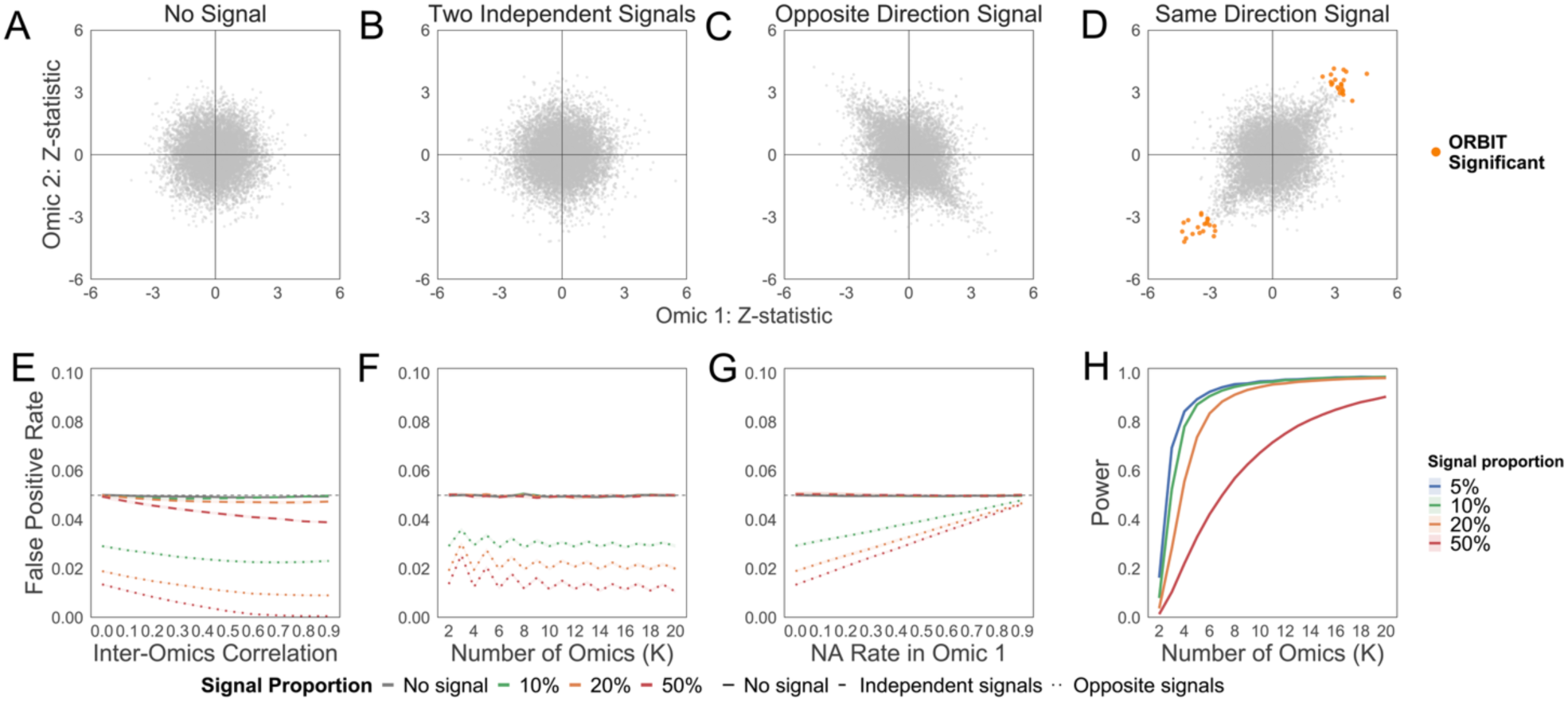
ORBIT remains well calibrated and detects concordant signals. (**A-D**) Four simulation scenarios for ORBIT in two omics. **(A)** No signals. **(B)** Two independent signals that have no correlations. **(C)** Correlated signals with opposite directions. **(D)** Concordant signals. **(E-G)** False-positive-rate at four proportions of signals (0%, 10%, 20%, 50%) for no signals (solid line), independent signals (dashed line) and opposite signals (dotted line) scenarios. **(E)** False-positive-rate under different between-omics correlation. **(F)** False-positive-rate with increasing numbers of omics layers. **(G)** False-positive-rate under different levels of feature missingness. **(H)** Power of ORBIT to recover concordant signal as a function of the number of omics layers, at four proportions of signals (5%, 10%, 20%, 50%).

We next assessed false-positive rate under three simulation settings. We simulated 10,000 features across two omics layers, with feature-specific Z-statistics drawn from a normal distribution. In each setting, we considered three scenarios, including no signals, independent signals and opposite signals. First, to reflect background correlation between omics layers in the absence of shared biological signal due to batch effects or overlapping samples, we varied the null correlation and found that ORBIT remained well calibrated (**Fig. 2E**). Second, as we increased the number of omics layers from 2 to 20 under independence, ORBIT maintained false-positive rate at or below the nominal level of 0.05, showing that it scales well as more layers are added (**Fig. 2F**). Third, to reflect differences in feature coverage across omics layers, such as between transcriptome and proteome, we randomly removed features from one layer. ORBIT remained well calibrated even when only 10% of features overlapped between the two omics (**Fig. 2G**). Finally, we evaluated ORBIT’s power to recover concordant signals. As the number of omics layers increased from 2 to 20, power increased steadily across concordant signal proportions 5%, 10%, 20% and 50%, showing increased detection of signals that were consistently ranked in the same direction across layers **(Fig. 2H).**

We further benchmarked ORBIT against RankProd, a related rank-based method [11]. Under the same simulation settings, we observed that RankProd’s false-positive rate increased with between-omics correlation, direction discordance and feature missingness, whereas ORBIT remained well calibrated (**Supplemental Fig. 3A-D**). Unlike ORBIT, RankProd ranks features separately in the up– and downregulated directions, allowing directionally discordant signals across layers to receive high rankings **(Supplemental Fig. 3E)**. Under independent, concordant and fully observed conditions, we observed comparable power between two methods (**Supplemental Fig. 3F**). Together, these results show that ORBIT maintains false-positive-rate control under heterogeneous multi-omics settings.

### ORBIT identifies coherent kidney injury programs across transcriptome and proteome

To evaluate ORBIT in real datasets, we first integrated transcriptomic and proteomic data from CKD versus healthy controls in human dissected tubulointerstitial tissue. We obtained proteomic data from the KPMP cohort, which quantified 6,555 proteins by LC-MS/MS [15], and transcriptomic data from the CPROBE cohort, which included 12,031 protein-coding transcripts by microarray [16] **(Fig. 3A)**. To demonstrate how ORBIT integrates different cross-omics patterns, we highlighted three representative genes **(Fig. 3B)**. Complement 3 (*C3*), which has been established as upregulated in CKD tubulointerstitial tissue [17], illustrates concordant upregulation across both omics and therefore ranked highly after ORBIT integration. Tissue inhibitor of metalloproteinases 3 (*TIMP3*) ranked highly in both omics but in opposite directions [18], resulting in a reduced integrated score in ORBIT. Finally, myosin heavy chain 7 (*MYH7*) showed little evidence in either layer and remained non-significant under ORBIT. These examples illustrate how ORBIT promotes concordant signals while down-weighting conflicting ones.

**Fig. 3.**
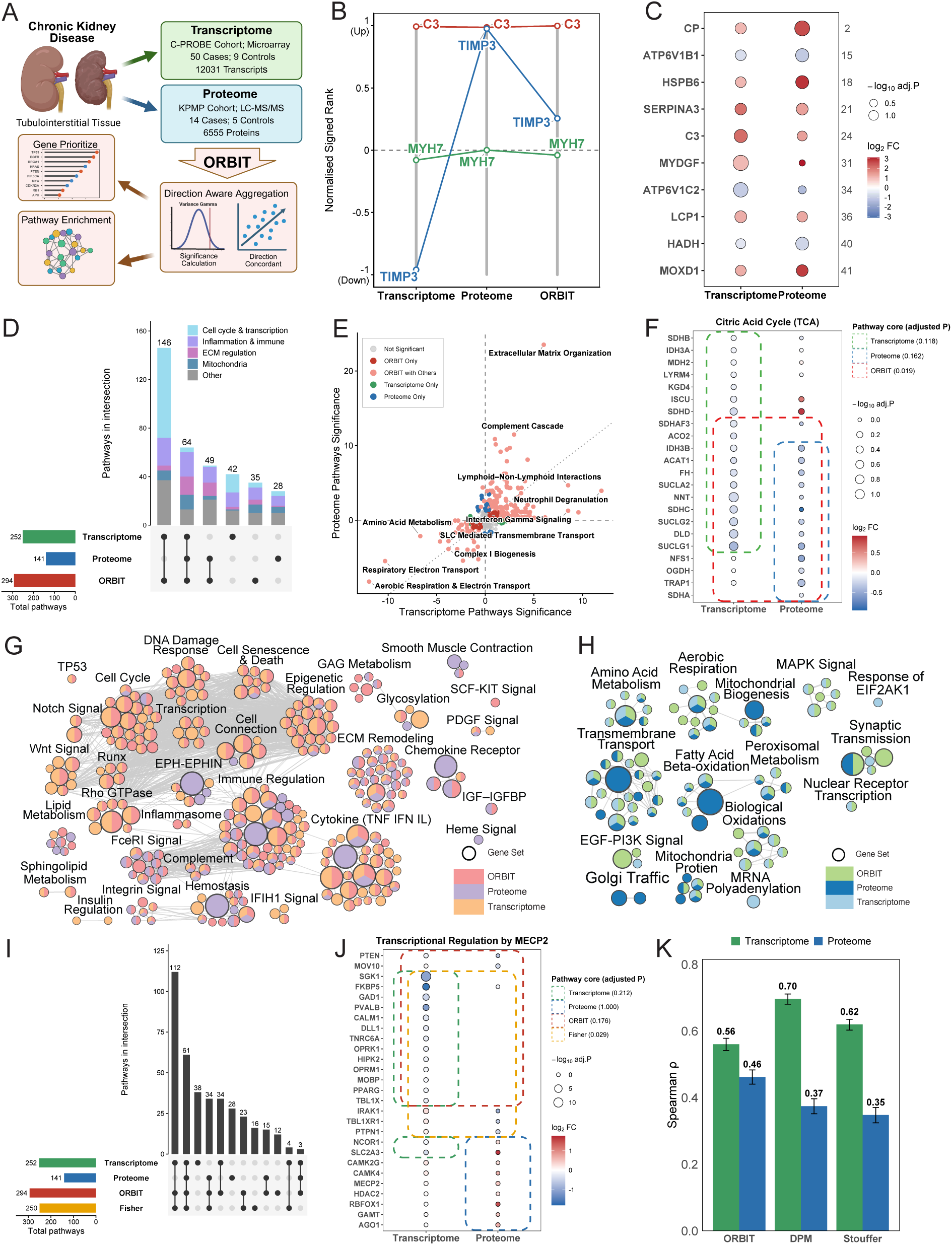
ORBIT integration of the transcriptome and proteome in chronic kidney disease. **(A)** Flowchart created in BioRender. Qiu, Z. (2026) https://BioRender.com/qjhgq9i. Transcriptomic and proteomic profiles of tubulointerstitial tissue were drawn from two independent cohorts: C-PROBE microarray data (50 cases, 9 controls, 12,031 transcripts) and KPMP LC-MS/MS proteomics (14 cases, 5 controls, 6,555 proteins). **(B)** Integrated statistic for three representative genes, shown as the normalized signed rank within each layer and after integration (sign*[1-rank/(m+1)]; m, number of features in each omics). **(C)** Top 10 ORBIT-ranked genes shared by both layers, shown separately for each layer. Numbers on the right indicate each gene’s rank in the full ORBIT gene list. **(D)** Significant pathways (adjusted P < 0.05) identified by GSEA against Reactome for each omics layer and ORBIT. Stacked bars are coloured by functional category. **(E)** Pathway-level comparison between the two omics layers. Each gene set is positioned by its signed significance, sign(NES)×−log_10_(adjusted P), in each layer and coloured by layer significance. The five most up– and downregulated pathways are labelled. **(F)** The REACTOME_CITRIC_ACID_CYCLE_TCA_CYCLE as an example of a pathway detected only after ORBIT integration. Enrichment maps of significant (adjusted P value < 0.05) upregulated (**G**) and downregulated (**H**) pathways under each omics layer and ORBIT. Each node is a gene set, sized by number of genes and connected to others by gene overlap. Node clusters were identified and labelled with AutoAnnotate [29]. (**I**) Significant pathways by GSEA for each omics layer, ORBIT and Fisher’s method, with direction for Fisher’s method taken from the sign of the mean log_2_(fold change). (**J**) REACTOME_TRANSCRIPTIONAL_REGULATION_BY_MECP2, a pathway called significant by Fisher’s method but not by ORBIT or either layer alone. **(K)** Spearman rank-correlation between the merged non-directional P values of each method (ORBIT, DPM, Stouffer) and the transcriptomic and proteomic P values across genes measured in both layers. Error bars denote 95% confidence intervals from bootstrap. In **C, F, J,** genes are coloured by log_2_(fold change) and point size indicates –log_10_(adjusted P value); dashed outlines mark leading-edge genes.

Among the top ten genes with the strongest concordant signals across the two omics layers, ORBIT ranked ceruloplasmin (*CP*) as the most upregulated and ATPase H^+^ transporting V1 subunit B1 (*ATP6V1B1*) as the most downregulated **(Fig. 3C)**. These findings are consistent with previous studies, as *CP* is a kidney-derived acute-phase protein reported to be upregulated in kidney tissue of CKD and an early urinary biomarker[19], whereas *ATP6V1B1* is well recognized as an intercalated-cell marker whose loss-of-function variants cause distal renal tubular acidosis with sensorineural deafness [20]. The marked downregulation of *ATP6V1B1* may therefore reflect loss or dysfunction of intercalated cells, which could contribute to the acidosis of late-stage CKD, consistent with its established role as a cell-type marker [21].

To identify the biological processes supported by ORBIT, we performed pathway enrichment analysis using GSEA and compared the resulting Reactome pathways [22] with those from the transcriptome and proteome analyzed separately. ORBIT recovered all 64 pathways shared between the proteome and transcriptome and identified 35 additional pathways that were not significant in either single omics layer **(Fig. 3D)**. At the pathway-level, ORBIT recovered a coherent CKD program characterized by upregulation of extracellular matrix (ECM) organization and the complement cascade and downregulation of respiratory electron transport, consistent with fibrotic and metabolic remodeling in diseased tubulointerstitial tissue [17, 23, 24] **(Fig. 3E)**. To illustrate how ORBIT identified pathways that did not reach significance in either individual layer, we used the citric acid cycle, a well-established biological process, as an example [25]. Although its core-enrichment genes were represented in both omics’ layers, neither layer alone reached significance (adjusted P value = 0.118, 0.162, in transcriptome and proteome respectively). In contrast, by prioritizing concordant genes across layers, ORBIT captured this pathway (adjusted P value = 0.019) **(Fig. 3F)**.

To summarize the biological processes captured by ORBIT and compare them with those seen in individual omics layer alone, we clustered the enriched pathways across ORBIT, the proteome and the transcriptome **(Fig. 3G, H, Supplemental Table 2)**. Upregulated pathways included ECM remodeling, cell cycle, immune regulation and cytokine signalling, whereas downregulated pathways included mitochondrial function and transmembrane transport. Notably, ORBIT prioritized up-regulated glycosaminoglycan (GAG) metabolism, which was not significant in either single-omics analysis, consistent with finding that FGF23 induces GAG accumulation to remodel the vascular extracellular matrix in CKD [26]. Together, these clusters are consistent with our current understanding of CKD pathophysiology [27].

To compare ORBIT with a conventional P value merging approach, we repeated the pathway analysis using Fisher’s method. Since Fisher’s method does not preserve direction, we assigned each feature the sign of its mean Log_2_FC across the omics layers to enable comparison [28]. We found that Fisher’s method was biased toward the transcriptome, likely because transcriptomic studies had larger sample sizes (59 vs 19) and greater measurement sensitivity, which systematically produced smaller P values. It favored transcriptome-supported features, missed 15 proteome-supported pathways which were covered by ORBIT, and failed to recover three pathways that were significant in both omics’ layers. In contrast, ORBIT recovered all pathways shared by the transcriptome and proteome **(Fig. 3I)**. To illustrate how Fisher’s method can prioritize layer-biased genes, we examined its top significant genes and found four were absent from the ORBIT top ten. Among these, *PDK4* and *LRATD2* showed opposite directions of effect between the two omics layers, whereas *RETREG1* and *CAVIN2* were strongly biased toward the transcriptome **(Supplemental Fig. 4)**. This shows that Fisher’s method can rank genes highly even when they are discordant or dominated by one layer rather than concordant across both. We also observed 16 pathways that were significant only using Fisher’s method. These pathways likely reflect Fisher’s tendency to prioritize genes with the smallest P values even when their effects were discordant. As an example, *MECP2*-regulated transcription was significant using Fisher’s method but not ORBIT or either individual omics layer. Its core enrichment revealed genes with opposite directions of effect and genes measured only in the transcriptome, indicating that the pathway was not supported by consistent cross-omic evidence **(Fig. 3J)**.

To compare ORBIT with other direction-aware multi-omics integration methods, we used the directional P-value merging (DPM) and Stouffer’s methods to combine the transcriptomic and proteomic results [8]. We compared how each method weighted the two omics layers by computing the Spearman rank-correlation between each method’s merged non-directional P values and the transcriptomic and proteomic P values across genes measured in both layers. ORBIT showed the smallest difference in correlation between the two omics layers, whereas DPM and Stouffer’s method correlated more strongly with the transcriptome than with the proteomic results **(Fig. 3K)**. Consistent with the simulation results, comparison with RankProd showed that genes with opposing directions between the transcriptomic and proteomic layers could still receive high RankProd rankings, whereas these genes were downweighted by ORBIT (**Supplemental Fig. 5**).

### ORBIT identifies robust transcriptomic programs in dilated cardiomyopathy

To further demonstrate the utility of ORBIT, we integrated ten transcriptomic studies of DCM heart tissues **(Supplemental Table 1, Fig. 4A)** [30–39]. We identified 1,299 significant genes measured in at least half of the studies with an ORBIT FDR-adjusted P value < 0.05. The top most significantly upregulated (*HTRA1*, *HMGN2* and *ASPN*) and downregulated (*SERPINA3*, *MYH6* and *HMOX2*) genes have all been previously reported in DCM transcriptome studies [28, 40, 41] (**Fig. 4B**). To ensure no single study dominated the overall result, we performed leave-one-study-out sensitivity analyses. Spearman correlation between the gene ranks across analyses remained stable across analyses **(Fig. 4C)**. In total, we identified 601 genes that were consistently identified across all leave-one-study-out analyses, forming a robust core DCM gene list **(Supplemental Table 4, Fig. 4D)**.

**Fig. 4.**
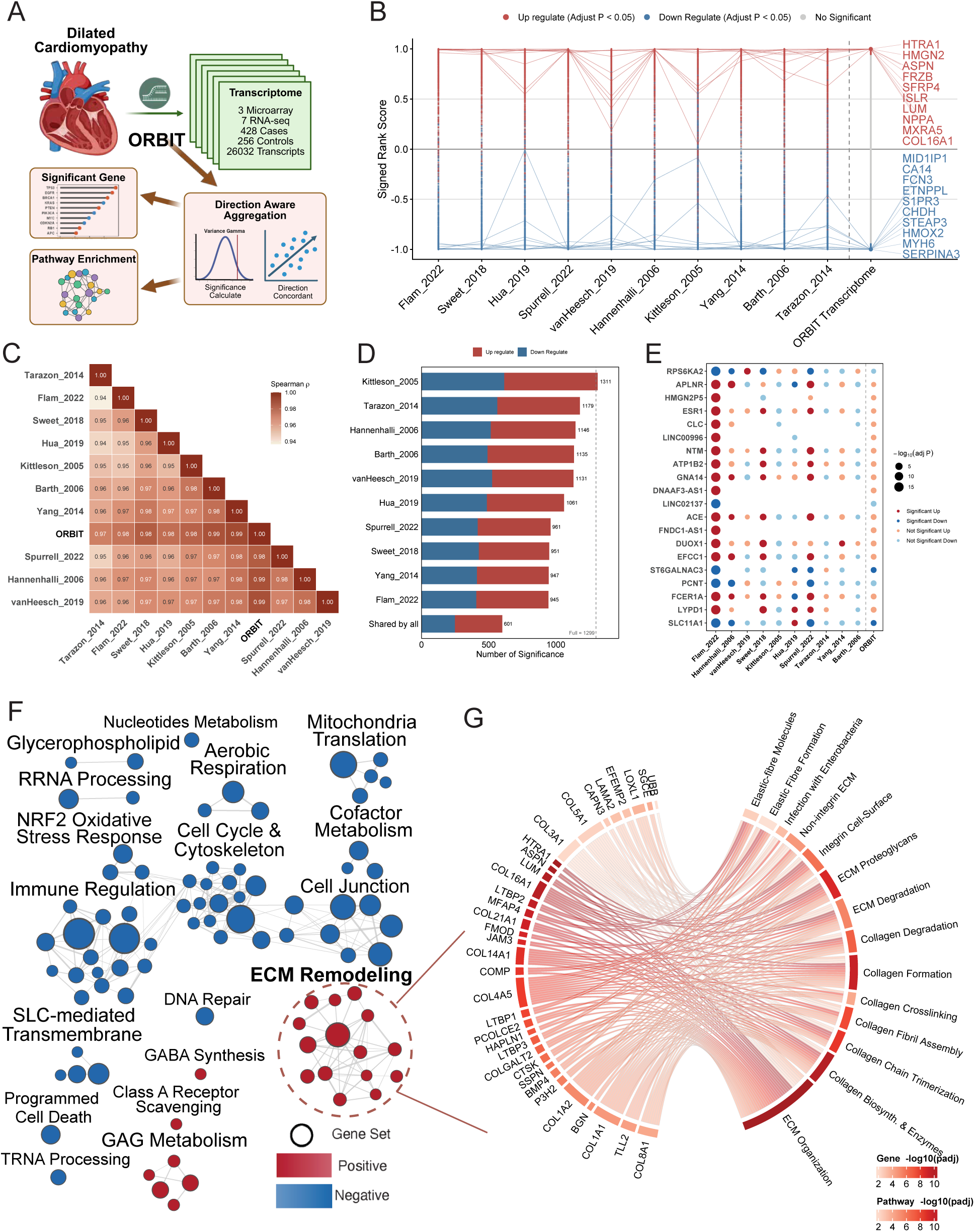
ORBIT integration of the transcriptome studies in dilated cardiomyopathy. **(A)** Flowchart created in BioRender. Qiu, Z. (2026) https://BioRender.com/n8zcepx. Transcriptomic profiles were drawn from ten independent studies of left ventricular tissue (428 cases, 256 controls, 26,032 transcripts). **(B)** Signed rank score of each gene in every contributing study and after integration across the transcriptomic layer. Points are colored by ORBIT significance, with upregulated genes in red, downregulated genes in blue. Lines trace the ten genes ranked highest and the ten ranked lowest by ORBIT, which are labelled on the right. Signed rank score is sign*[1-rank/(m+1)]. **(C)** Spearman correlation between leave-one-out integrations and the full ORBIT ranking. **(D)** Number of significant genes per leave-one-out studies, split by direction. “Shared by all” indicates the number of significant genes shared among all leave-one-out analyses; and the dashed line marks the number of significances using all studies. **(E)** Genes in the top 5% of the Fisher’s method ranking that fall furthest below it under ORBIT, showing genes prioritized by Fisher’s method but not by ORBIT. Colour indicates sign of log_2_(Fold Change) together with significance and point size the per-omic –log_10_(adjusted P value). **(F)** Enrichment maps of significant (adjusted P value < 0.05) up and downregulated pathways by ORBIT. Each node is a gene set with color represents direction, size represents number of genes in the pathway. Nodes are connected by gene overlap. Node clusters were identified and labelled with AutoAnnotate [29]. **(G)** Chord diagram of the extracellular matrix pathway clusters significant by GSEA and their core-enrichment genes which are also the robust significant genes. Ribbon and segment color represents the –log_10_(adjusted P value) of the gene and the pathway respectively.

To compare gene prioritization between ORBIT and Fisher’s method on the gene discovery, we selected the top 5% of genes ranked by Fisher’s method and ranked them by the difference between their Fisher and ORBIT ranks. We highlighted genes such as *RPS6KA2* and *APLNR* that Fisher’s method prioritized despite directionally discordant effects or strong dependence on individual studies through non-directional P value merging **(Fig. 4E)**.

To further characterize the biological mechanisms underlying DCM gene expression, we conducted GSEA pathway analysis and clustered the Reactome pathways by direction of change **(Fig. 4F)**. The transcriptome profile of DCM was characterized by upregulated ECM and GAG metabolism and downregulated innate immune regulation, cell junction, cell cycle and mitochondrial functions, aligning with established DCM biology at transcriptome level [40, 42, 43]. We then asked whether the strongest robust genes pointed to the same biology. We performed Reactome over-representation analysis (ORA) on the 601 robust significant genes and compared the resulting pathway significance against that from GSEA. The enriched pathways converged on an upregulated ECM regulating cluster **(Supplemental Table 5, Supplemental Fig. 6),** indicating that ECM remodeling is supported by both the top genes and the broader rank structure. To identify the genes driving this signal, we linked the ECM-cluster pathways to their core-enrichment genes **(Fig. 4G)**. The core-enrichment genes also included established extracellular matrix components, including *COL1A1*, *COL1A2*, *COL3A1*, *LUM* and *FMOD*, several of which have previously been associated with myocardial fibrosis and identified as reproducible transcriptional changes in DCM [44]. Together, these genes support a coordinated programme of collagen deposition and extracellular matrix remodeling in DCM and other cardiomyopathies [45].

### ORBIT identifies concordant regulation across transcriptome, translatome and proteome in dilated cardiomyopathy

To further explore the applications of ORBIT, we used it to characterize concordant biological processes of DCM across the transcriptional, translational and protein levels. To avoid bias toward the transcriptomic layer due to its large number of contributing studies, we used the transcriptomic summary statistics from the above ORBIT integration result. We derived the translatome, expressed as translation efficiency, from the available cardiac translatome study [39]. We assembled the proteome from four published, non-overlapping DCM proteomic studies [46–49] **(Supplemental Table 1, Fig. 5A)** and integrated them using ORBIT to obtain a single proteomic summary **(Supplemental Fig. 7)**. We then integrated the three omics layers using ORBIT. Among the top-ranked genes, the downregulated genes reflected failing heart biology, including the α-to-β myosin switch (*MYH6*) [50], reduced macrophage markers (*CD163*) [51] and impaired calcium handling (*ATP2A2*) [52], whereas the upregulated genes pointed to ECM remodeling (*LUM, ASPN*) [53, 54] and fibrosis (*LTBP2, PTN*) [55, 56] (**Fig. 5B**).

**Fig. 5.**
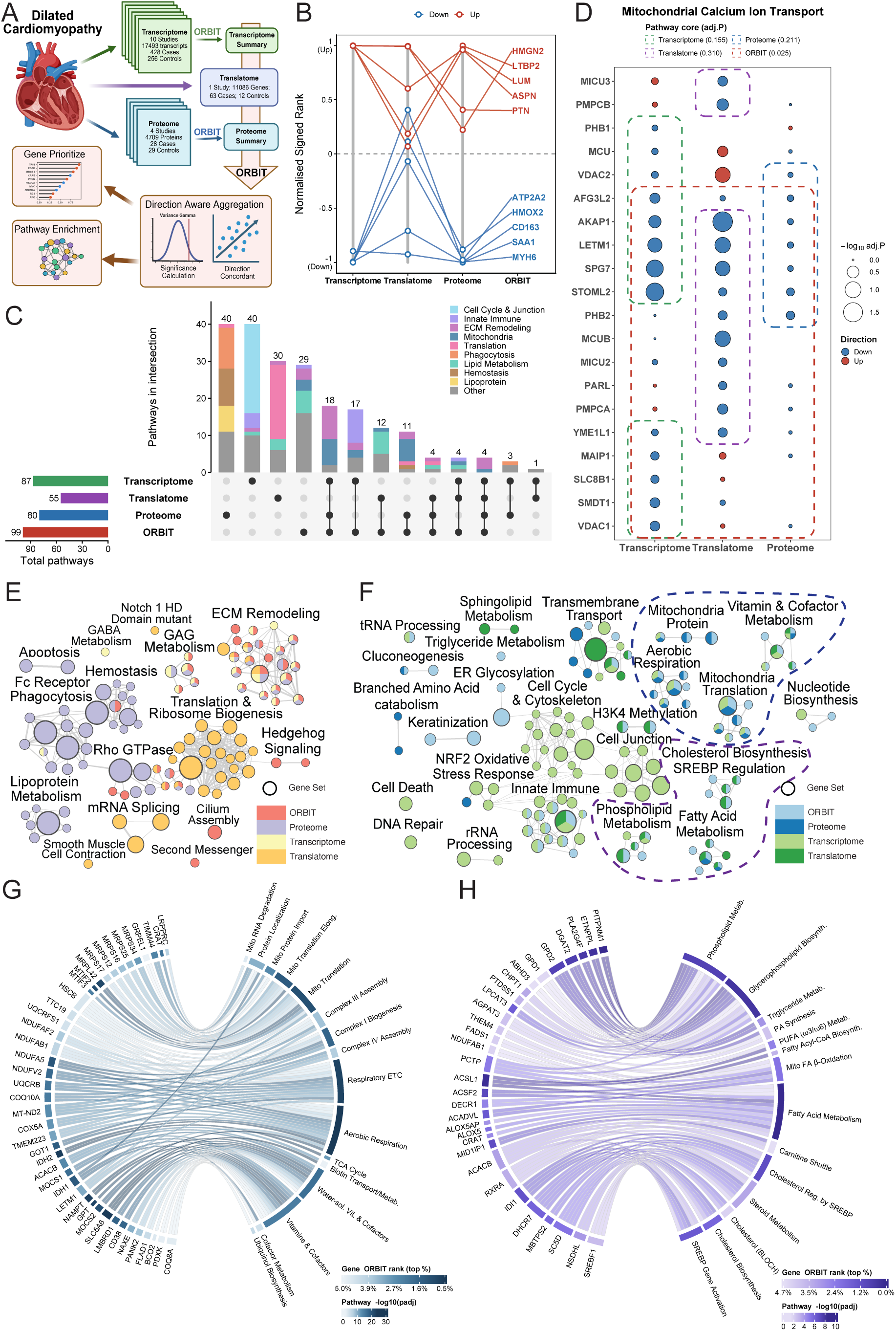
ORBIT integration of transcriptome translatome and proteome in dilated cardiomyopathy. (**A**) Flowchart created in BioRender. Qiu, Z. (2026) https://BioRender.com/yeyd62x. Three layers were integrated: transcriptomes from 10 studies (428 cases, 256 controls, 17,493 genes), a translatome from one ribosome-profiling study (63 cases, 12 controls, 11,086 genes), and proteomes from 4 studies (28 cases, 29 controls, 4,709 proteins). **(B)** Normalized signed rank of the five genes ranked highest and the five ranked lowest by ORBIT, shown within each layer and after integration. Normalized signed rank is sign*[1-rank/(m+1)]. **(C)** Significant pathway (Adjusted P value < 0.05) by GSEA against Reactome under each omics layer layers and ORBIT, stacked bars are coloured by functional category. **(D)** The REACTOME_MITOCHONDRIAL_CALCIUM_ION_TRANSPORT as an example of a pathway detected only after integration. Genes are shown with log_2_(Fold Change) by colour and per-omic –log_10_(adjusted P value) by point size; dashed outlines mark the leading-edge genes driving enrichment in each layer. **(E)** Enrichment maps of significant (adjusted P value < 0.05) up regulated pathways under each omics layer and ORBIT. Each node is a gene set, sized by number of genes and connected to others by gene overlap Node clusters were identified and labelled with AutoAnnotate [29]. **(F)** Enrichment maps of significant (adjusted P value < 0.05) down regulated pathways. **(G)** Chord diagram of the mitochondria and cofactor metabolism clusters significant by GSEA and their core-enrichment genes which are also the robust significant genes. Ribbon colour and segment colour give the –log_10_(adjusted P value) of the gene and the pathway respectively. **(H)** Chord diagram of the lipid and fatty acid metabolism clusters significant by GSEA.

To identify the underlying biological processes supported across the three omics layers, we performed GSEA. We observed little agreement among the individual layers, whereas ORBIT recovered all four overlapping pathways shared across layers and identified 29 additional pathways that were not significant in any individual layer **(Fig. 5C)**. As an example of the ORBIT-specific pathways, we examined the mitochondrial calcium ion, as impaired mitochondrial calcium homeostasis is a recognised feature of DCM [57]. Its core-enrichment genes demonstrated that ORBIT could aggregate weak and incomplete signals from different omics layers to identify relevant biological processes **(Fig. 5D)**. Further clustering of the significant pathways showed that ORBIT captured coherent programs, including upregulated ECM remodeling and glycosaminoglycan metabolism, and downregulated mitochondrial and cofactor metabolism together with lipid and fatty-acid metabolism **(Fig. 5E, F, Supplemental Table 3)**. ECM remodeling had already been identified at the transcriptome level **(Fig. 4F)**. In addition, omics integration further revealed the downregulated mitochondria, cofactor, and lipid-related pathway cluster, which converged on a coherent picture of impaired cardiac energy metabolism, a recognised hallmark of the failing heart [58].

To identify the genes driving these signals, we generated chord diagrams for the mitochondria and cofactor clusters and highlighted the core-enrichment genes ranked in the top 5% of the ORBIT list **(Fig. 5G).** The leading-edge genes included multiple components of the mitochondrial respiratory chain, including *NDUFV2*, a core subunit of complex I [59], and *COX5A*, a subunit of complex IV [60], consistent with the coordinated impairment of mitochondrial respiratory pathways in DCM. Similarly, within the lipid and fatty-acid metabolism clusters, *ACSL1*, a key enzyme involved in fatty-acid activation and oxidation and previously reported as a key gene in DCM, ranked near the top of the ORBIT list (rank = 80/18,220) [61]. *NDUFAB1*, a component of mitochondrial respiratory chain complex I, appeared in the leading edge of both clusters and ranked within the top 5% of all genes (rank = 817/18,220), and has been shown to support cardiac mitochondrial bioenergetics and function [62] **(Fig. 5H)**.

## Discussion

In this study, we introduce ORBIT, a rank-based statistical framework for multi-omics integration that jointly models the rank and the direction of effect of each feature across heterogeneous datasets. ORBIT addresses a setting in which conventional meta-analysis is particularly difficult such as when datasets differ in assay technology, sample size, statistical model and feature coverage. We showed via simulations that ORBIT remains robust to between-omics correlation, high dimensionality and missingness. Across CKD and DCM, ORBIT recovered reproducible biological programmes supported consistently across molecular layers. An important distinction between ORBIT and conventional significance-combination approaches is the hypothesis being tested. Methods such as Fisher’s combined probability test ask whether there is aggregate evidence against the null across datasets, but do not require the underlying effects to agree in direction. Directional P value approaches, including DPM and directional Stouffer-type procedures, incorporate directional information into the merging step, but their resulting significance remains dependent on the calibration and magnitude of the input P values or test statistics [8]. ORBIT instead asks whether a feature repeatedly occupies an extreme rank in consistent direction, such that opposing signals cancel rather than reinforce one another.

Rank transformation provides a practical advantage when integrating heterogeneous assays. A P value reflects both the strength of evidence and estimation precision, and its absolute scale can therefore differ substantially across studies with different sample sizes, assay sensitivities and statistical models. Rank-based methods do not eliminate these differences in power. Instead, they replace absolute significance values with within-dataset ranks, reducing sensitivity to differences in statistical scale at the cost of discarding information about the magnitude of evidence. ORBIT extends this principle by incorporating direction, background correlation and incomplete coverage. Consistent with this design, ORBIT remained well calibrated under these sources of heterogeneity in simulations and showed a more balanced relationship with the transcriptomic and proteomic layers than significance-combination methods in the CKD analysis (**Fig. 3K**, **Supplemental Fig. 4).**

Prioritizing concordance can reduce the influence of layer-specific technical or compositional effects that might otherwise dominate integration. For example, the upregulation of lipoprotein and hemostasis pathways seen only in the proteome may reflect blood carryover rather than tissue biology in the DCM analysis. Similarly, translation– and ribosome-related pathways, including eukaryotic translation elongation, were detected only in the translatome, reflecting biologically meaningful omics-specific signals that were not supported across the other omics layers.

Conversely, ORBIT highlighted biological processes for which concordance evidence was present across datasets but did not reach significance in individual omics layers. These included ECM and inflammatory together with impaired mitochondrial and metabolic function in CKD, both consistent with known disease biology [27]. The citric-acid-cycle pathway in kidney tissue illustrates this principle, while the multi-omics DCM analysis similarly strengthened evidence for impaired mitochondrial, cofactor and lipid metabolism and prioritized genes include *NDUFV2, COX5A,* and *NDUFAB1*. Thus, pathways detected only after ORBIT integration reflect concordant evidence that is incompletely captured by individual datasets rather than signals created by the integration procedure. Our method has several limitations. First, ORBIT operates on summary-level statistics and therefore cannot model individual-level covariance or identify patient-specific multi-omic states. Second, by relying on ranks, ORBIT removes the magnitude of evidence and compresses differences in statistical evidence within each study. This reflects a deliberate trade-off favouring consistency, because ORBIT does not accumulate study-level precision for association testing in the manner of conventional meta-analysis. To provide an alternative when retaining the original statistical evidence is desirable, we implemented ORBIT-P as an extension within the released ORBIT software. ORBIT-P adopts the same statistical model but retains the original P values, providing a directional extension of Fisher’s method. We further demonstrate that ORBIT-P maintains a well-calibrated false positive rate under inter-omic correlation, multi-layer designs, and missingness, while retaining higher power than DPM in our simulations, owing to its precise modeling of both effect direction and P-values **(Supplemental Fig. 8).** Third, cross-omic integration requires features from different molecular layers to be mapped onto a common feature space. In our analyses, transcriptomic and proteomic features were mapped to genes, but such mappings are not always one-to-one and may collapse transcript-, isoform– or protein-specific signals. This limitation becomes more important for molecular layers such as metabolomics or epigenomics, where mapping features to genes can be indirect or ambiguous. Fourth, ORBIT is designed to prioritize concordant signals, meaning that strongly discordant effects across molecular layers will receive reduced integrated evidence even when such divergence reflects genuine biology, for example through post-transcriptional or other layer-specific regulatory process [63].

Characterization of such signals therefore require complementary analysis of the individual omics layers. Fifth, the variance-gamma null distribution provides a continuous approximation to the finite discrete rank distribution and is consequently conservative, particularly when the number of ranked features is small [64]. Finally, ORBIT requires estimation of background correlation between datasets. This can be obtained by permutation when individual-level data are available, but otherwise must be approximated from non-significant features, which may contain residual biological signal and make the test conservative.

ORBIT is complementary rather than universally preferable to existing meta-analytic and multi-omics approaches. When comparable effect estimates and uncertainties are available from similarly designed studies, conventional fixed– or random-effects meta-analysis retains quantitative information that rank transformation deliberately discards. ORBIT is instead intended for settings in which datasets differ substantially in assay technology, statistical pipeline or feature coverage, and only summary-level evidence is available. Its minimal input requirements also make the framework readily extendable to other applications, including integration of eQTL, pQTL and metabolite-QTL evidence, pseudobulk signals from single-cell studies, and additional molecular layers such as metabolomic or epigenomic data.

## Methods

### ORBIT statistical framework

For each feature i in dataset j, let *r_i_*_j_ ∈ {1, …, *m*_j_} denote its within-dataset rank and 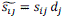 its aligned direction of effect (*s_i_*_j_ ∈ {−1, +1} that denotes the feature-level sign and *d*_j_ ∈ {−1, +1} that denotes a dataset-level reference orientation). Let 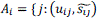 is observed} denote the set of datasets in which the feature is measured, and *k_i_* = |*A_i_*| denote the number of contributing datasets. ORBIT defines the directional integrated statistic as: 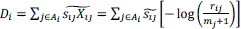. Under the null, each signed score is distributed as Laplace(0,1), and assuming independence between contributing datasets, *D_i_* follows a symmetric variance-gamma (VG) distribution [65] or the symmetric generalized Laplace distribution[66]. Here we used the general four-parameter VG parameterization (σ, ν, θ, µ) of Madan et al. [13], with ν*_i_* = 1/*k_i_*, 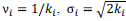, θ = 0, and µ = 0. The number of contributing datasets thus determines the shape of the null, and its scale the dispersion of the integrated statistic (derivation in **Supplemental Note**).

The VG null derivation assumes that the signed rank scores contributed by different datasets are independent under the null. In practice, datasets may be correlated owing to sample overlap, batch effects, or coupled normalization, which inflates the variance of the integrated statistic *D_i_* even for features carrying no true signal. ORBIT therefore applies a Brown-style correction that rescales the VG null parameters using an estimate of background between-dataset correlation [7]. This correlation is estimated from a permutation null when individual-level data are available, or otherwise from features showing no evidence of association in any contributing dataset. Details of the estimation procedure and parameter adjustment are provided in **Supplemental Note.** The VG parameters are therefore 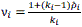, and 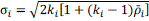.

Existing implementations of the VG distribution are not designed for the extreme tail probabilities required in omics analyses; the implementation used in [67] returns probabilities below 10⁻¹⁰ as zero and provides limited accuracy for P < 10⁻⁶. ORBIT therefore implements a dedicated log-space evaluator that combines closed-form solutions for special cases of ν*_i_* with Gauss-Laguerre quadrature for the general case, with all computations performed in log space to avoid underflow. We validated the implementation against an importance-sampling Monte Carlo reference (tail probabilities down to 10⁻¹⁴) and benchmarked it against the existing VG package. Our analytical values agree with the empirical tail probabilities to within sampling error, whereas the package deviates progressively in the tail (**Supplemental Fig. 9)**. Finite log P_i_ values are returned for *d_i_* up to at least 5,000 (log P_i_ ≈ −2400) far beyond any value encountered in multi-omics applications. Full details of the evaluator are provided in **Supplemental Note**.

ORBIT-P is an extension of ORBIT that uses the original per-omic P values in place of rank-transformed values. Under the null, a valid P value is uniformly distributed on (0, 1), so the same signed transformation and integration framework applies directly to the input values, giving 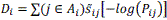. For fully directional inputs, this statistic is equivalent to the DPM statistic

[8]. DPM evaluates this signed sum against a scaled χ² distribution calibrated by the empirical Brown’s method [8], whereas ORBIT-P evaluates it against the exact variance-gamma null derived above, which accounts for its higher power in simulations **(Supplementary Fig. 8)**.

### Data Simulation

ORBIT distribution is validated through two simulations. For two omics layers, the exact discrete-rank null was obtained by enumeration of all rank pairs and sign combinations; for three to five layers, the discrete null was estimated from 10⁹ rank permutations. Simulations covered 2,000, 5,000, 10,000, and 20,000 ranked features per dataset. To assess the adequacy of the VG approximation itself, we simulated signed sums of 2^−10^ independent uniform scores and compared analytical VG tail probabilities with Monte Carlo estimates.

ORBIT’s performance was evaluated through simulations. For *N* features measured across *K* omics, each feature’s vector of Z-statistics was drawn from a multivariate normal distribution, N(**μ**, Σ), where Σ is an equicorrelated covariance matrix. For null features, the common correlation ρ_null_, representing background correlation between omics datasets. For signal features, the common correlation was set to ρ_signal_. Signal magnitude and direction were specified through the mean vector **μ**.

Non-signal features carried µ = 0, whereas signal features carried a per-omic mean of magnitude µ_signal_ = 1.5. We simulated three types of signals. The first was a concordant signal, in which a feature shared the same direction across all omics (**μ** = ±µ_signal_**1_K_**, ρ_null_ = 0, ρ_signal_=0.95). The second was a discordant signal, illustrated with two omics, in which a feature had µ = + 1.5 in one omics and µ = − 1.5 in the other. The third comprised two independent signals, in which each of the two omics carried signal independently of the other, so that a feature could carry signal in one omic, the other, or both. In every case, ORBIT operated on the per-omic signs of *Z* and the statistic ranks, whereas the comparators (Fisher’s, Brown’s, and Stouffer’s) combined the two-sided *P*-values P = 2Φ (−|*Z*|).False positive rate was assessed under three departures from the assumptions of independence and complete observation: (i) between-omics correlation, with ρ increased from 0 to 0.9 in steps of 0.1; (ii) dimensionality, with k taking every integer value from 2 to 20 under independence; and (iii) missingness, with features randomly omitted from one omics at rates of 0-90% in steps of 10%. Each grid point was replicated 20 times with independent seeds, each replicate comprising 10 000 features. False positive rate was defined as the proportion of null features with unadjusted P < 0.05, against a nominal level of 0.05; no multiplicity correction was applied. Curves show the mean across replicates with 95% confidence intervals. ORBIT was evaluated under three null scenarios: no signal, independent signals, and opposite signals. In the opposite-signal scenario, signs split as evenly as possible between positive and negative among K omics (K/2 each for even K; the closest split, chosen at random, for odd K). Additionally, ORBIT-P was evaluated under the no-signal scenario against four comparators: Fisher’s combined probability test, directional Stouffer’s method, DPM and Brown’s method as implemented in the empirical Brown’s method **(Supplemental Fig. 8).** To evaluate power, we simulated 10,000 features with equal numbers of up– and downregulated signal features. Signal features had a common concordant mean effect µ_signal_ = 3. Correlation was set to ρ_null_ = 0 among null features and ρ_signal_ = 0.9 among signal features, so that true signals covary strongly across omics while the background remains independent. Signal proportions of 5%, 10%, 20%, and 50% were evaluated as K increased over every integer from 2 to 20, with 20 replicates per grid point. Power was defined as the proportion of true signal features recovered at a Benjamini-Hochberg adjusted P < 0.05.

### Data sources and processing

ORBIT input requires summary-level input data. For CKD transcriptome, we analysed microarray data obtained from C-PROBE cohort (GSE180395) [16]. We grouped samples by histological diagnosis into healthy controls (living donors and unaffected tissue from tumour nephrectomies) and CKD subtypes (lupus nephritis, Focal Segmental Glomerulosclerosis, Focal Global Glomerulosclerosis, diabetic nephropathy, membranous nephropathy, and two composite categories covering the remaining glomerular and non-glomerular diseases). We assessed differential expression using limma [68] under a cell-means design with one coefficient per subtype, contrasting the unweighted average of the CKD subtypes against the unweighted average of the two healthy groups so that each subtype contributes equally irrespective of sample size. We annotated genes with `org.Hs.eg.db` and restricted the analysis to protein-coding genes with a valid HGNC symbol. Where multiple identifiers mapped to the same HGNC gene symbol, we retained the one with lowest P value.

For the DCM transcriptome, we included ten non-overlapping studies from a recent systematic compilation [30–39], restricted to non-failing and DCM samples. We excluded donors under 18 years and adjusted for age and sex where available. We included only studies with at least five cases and five controls (**Supplemental Table 1**) and assessed differential expression using limma[68]. We retrieved summary-level CKD proteome results from the KPMP data portal [15] and DCM proteome and translatome results from the original publications [39, 46–49].

### ORBIT integration and sensitivity analysis

We estimated inter-omic correlation by permutation where individual-level data were available and from a signal-depleted background otherwise. For the DCM transcriptome, samples were permuted 1,000 times and the mean correlation across permutations was used. For the CKD analysis and the DCM multi-omic integration analyses, the original individual-level data were unavailable, so we estimated correlation from features that were non-significant in the contributing datasets. For the DCM transcriptome analysis, we assessed robustness using a leave-one-out analysis. We repeated the ORBIT integration after excluding each study in turn and compared the resulting gene rankings and significant genes with those from the complete analysis.

### Comparison with other integration methods

We compared ORBIT with Fisher’s method, RankProd, DPM and direction-aware Stouffer integration. For Fisher’s method, we combined the individual P values into the statistic X² = −2 Σ ln pᵢ and obtained the integrated P value from the corresponding chi-squared.

For RankProd, we used the RankProd package (v 3.36.0) [11]. We converted each per-omic P value and direction into a signed Z-statistic, 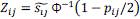, and supplied the resulting feature-by-omic matrix to the one-class `RankProducts` function. RankProducts ranks features separately in the up– and downregulated directions and returns direction-specific P values which we then combined into a two-sided integrated P value as min(2 ⋅ min(p^up^, p^down^), 1). To match ORBIT’s handling of missing data, we retained features observed in at least one omics layer.

We applied DPM and direction-aware Stouffer methods using the R package ActivePathways (v2.0.6) with a constraints vector of c(1, 1) [8]. Because ActivePathways couples P-value merging to its own ranked pathway-enrichment procedure and does not report direction-aware pathway-level output, a comparison with ORBIT at the pathway level was not possible. To place the methods on a comparable footing, we instead assessed their integration behaviour at the gene level. For genes measured in both omics’ layers, we computed the Spearman rank-correlation between each method’s merged (non-directional) P values and the transcriptomic and proteomic P values, respectively. We calculated 95% confidence intervals by non-parametric bootstrap, resampling genes with replacement 5,000 times and taking the 2.5th and 97.5th percentiles of the resulting bootstrap distribution.

### Pathway enrichment and plot

We performed pathway analysis principally by GSEA against Reactome[22], KEGG[69], GO:BP[70] and Hallmark [71], with Reactome used for the main figures and the remaining in supplemental materials. ORBIT, each single-omic input, and Fisher’s combined probability test were analysed under the same gene sets and settings for comparison. We ordered the pre-ranked list by the directional statistic sign(*D_i_*) × (− log_10_ *P_i_*). Although this differs from the original application of GSEA, signed statistics are widely used for pre-ranked GSEA and provided a common directional ranking across omics layers analyzed using different statistical methods. We ran GSEA using the `GSEA` function of the clusterProfiler R package (v4.18.4) [72], which implements the pre-ranked, gene-set-permutation algorithm of fgsea (v1.36.2) with the weighted scoring scheme. Gene sets were restricted to 10-500 members and the random seed was fixed for reproducibility. Enrichment was computed without filtering and significance applied afterwards at a Benjamini-Hochberg adjusted P-value. Enriched pathways were merged and visualised in Cytoscape (v3.10.4) using standard protocols [73].

We additionally performed over-representation analysis on the features called significant by ORBIT, partitioned them into up– and downregulated sets according to their integrated direction and tested each set separately. We assessed each set by hypergeometric test using the enricher function of `clusterProfiler` against the same 10-500 member gene sets, with the background universe defined as the complete set of features for which ORBIT returned an integrated result. We retained both raw and Benjamini-Hochberg adjusted P values. Reporting ORA alongside GSEA both characterizes the significant feature set directly and demonstrates that ORBIT output is available with threshold-based as well as rank-based pathway methods.

We produced all figures in R (4.5.2) using ggplot2, circlize, ComplexHeatmap, and patchwork, as well as Cytoscape (v3.10.4). The flowchart was created with BioRender.com.

## Data Availability

Transcriptome data are retrieved from the Gene Expression Omnibus (GEO), including CKD transcriptome data from the C-PROBE cohort (GSE180395), DCM transcriptome (GSE141910, GSE116250, GSE5406, GSE135055, GSE1869, GSE55296, GSE126569, GSE46224, GSE3585). Summary-level CKD proteome results were obtained from the Kidney Precision Medicine Project (KPMP) data portal (https://atlas.kpmp.org). DCM proteome, translatome and one transcriptome summary statistics were taken from the supplemental materials of the original publications.

## Code Availability

ORBIT is available as an R package at https://github.com/yang-luo-lab/ORBIT, with an interactive web version https://yang-luo-lab.github.io/Rank-based-integration-identifies-convergent-disease-mechanisms-across-omics/. Data processed and figure plotting code is available at https://github.com/yang-luo-lab/ORBIT-manuscript.

## Contributions

Z.Q., J.N. and Y.L. conceived the study. Z.Q. and Y.L. developed the ORBIT method and R package and performed the simulations. Z.Q., Y.L., D.P. and L.J. refined the methodology and code. Z.Q. and Y.L. analyzed the data. Z.Q., Y.L., J.N., A.L. and K.B. provided biological interpretation. Y.L. and J.N. supervised the work. all authors contributed to the review and editing of the manuscript.

