## Supplementary Notes for "Rank-based integration identifies convergent disease mechanisms across omics"

### Supplemental Notes

**ORBIT statistical framework**

ORBIT integrates summary-level statistics across multiple omics datasets to identify features that show consistent directional evidence across datasets. For each feature, ORBIT asks whether it tends to occupy an extreme rank in the same biological direction across the datasets in which it is measured. The method therefore evaluates cross-dataset directional concordance rather than the magnitude of association statistics. For dataset j, let $m_j$ denote the total number of features measured after feature harmonisation. For each feature $i$ in dataset j, let $r_{ij} \in \{1, \ldots, m_j\}$ denote its rank according to the chosen evidence statistics, with $r_{ij} = 1$ representing the strongest evidence. Ranks may be derived from any statistic that conveys significance, including P values and effect sizes.

We convert the within-dataset to a common scale using

$$u_{ij} = r_{ij}/(m_j + 1)$$

where the denominator $m_j + 1$ makes the transformed ranks lie strictly between 0 and 1 and thus makes their scale comparable across datasets with different number of measured features. Under the null hypothesis, ranks $u_{ij}$ are uniformly distributed across the $m_j$ possible rank positions. Consequently, the normalized ranks form a discrete uniform distribution on its rank grid. As $m_j$ increases, this discrete distribution converges to a continuous Uniform (0,1) distribution.

We then transform the normalized rank to a log scale,

$$X_{ij} = -\log u_{ij}.$$

Under the continuous-rank approximation, $u_{ij}$ ~Uniform (0,1) and consequently $X_{ij} \sim \mathrm{Exp}(1)$. This transformation means that more extreme ranks produce larger evidence scores, while placing ranks from datasets with different numbers of measured features on a common scale. To incorporate direction, we define direction parameters, $s_{ij} \in \{-1, +1\}$ that denotes the feature-level sign and $d_j \in \{-1, +1\}$ that denotes a dataset-level reference orientation. Their product, $\widetilde{s_{ij}} = s_{ij}\, d_j$, aligns the direction across datasets so that the same sign had a consistent biological interpretation in every layer. Under the null, the underlying directional test statistic is symmetric about zero and its sign, $\widetilde{s_{ij}}$, is independent of its rank, $X_{ij}$. The signed rank score $\widetilde{s_{ij}} X_{ij}$

therefore follows a symmetric distribution centered at zero. Under the continuous rank approximation, it follows a Laplace distribution with location 0 and scale 1, that is,

$$\widetilde{s_{ij}}X_{ij} \sim \text{Laplace}(0,1)$$

For each feature $i$, let $A_i = \{j: (u_{ij}, \widetilde{s_{ij}}) \text{ is observed}\}$ denote the set of datasets in which the feature is measured, and let $k_i = |A_i|$ denote the number of contributing datasets. ORBIT defines the directional integrated statistic as:

$$D_i = \sum_{j\in A_i} \widetilde{s_{ij}X_{ij}} = \sum_{j\in A_i} \widetilde{s_{ij}} \left[-\log\left(\frac{r_{ij}}{m_j+1}\right)\right],$$

which simultaneously encodes cross-dataset coherence and overall direction. This means that strongly ranked features have larger scores, weakly ranked features have scores closer to zero, and opposing directions produce scores with opposite signs.

Under the assumption that the contributing datasets are independence under the null, $D_i$ is a sum of $k_i$ i.i.d $\text{Laplace}(0,1)$ variables. By definition, the characteristic function of a Laplace (0,1) variable is

$$\varphi_{\text{Laplace}(0,1)}(t) = (1+t^2)^{-1}$$

Therefore,

$$\varphi_{D_i}(t) = (1+t^2)^{-k_i}.$$

This is the characteristic function of a symmetric variance-gamma (VG) distribution [67] or the symmetric generalized Laplace distribution[68]. Equivalently, the sum can be viewed as the difference between two independent Gamma variables. That is, $G_1, G_2 \sim \text{Gamma}(k_i, 1)$ with characteristic function $\varphi_{G_1-G_2}(t) = (1-it)^{-k_i}(1+it)^{-k_i} = (1+t^2)^{-k_i}$, which matches $\varphi_{D_i}$ exactly. The number of contributing datasets therefore determines the shape of the null distribution, while its scale determines the dispersion of the integrated statistic.

Here, we used the general four-parameter VG parameterization $(\sigma, \nu, \theta, \mu)$ of Madan et al. [12]

whose characteristic function is $e^{i\mu t}(1 - i\theta\nu t + \frac{1}{2}\sigma^2\nu t^2)^{-1/\nu}$. By matching this expression to $\varphi_{D_i}(t) = (1+t^2)^{-k_i}$ term by term, we define $\nu = 1/k_i$, $\sigma = \sqrt{2k_i}$, $\theta = 0$, and $\mu = 0$. The asymmetry $(\theta)$ and location $(\mu)$ parameters are 0 due to the symmetry of the Laplace scores under the null. That is,

$$D_i \sim \mathrm{VG}\left(\sqrt{2k_i}, 1/k_i, 0, 0\right),$$

and the variance follows as $\mathrm{Var}(D_i) = \sigma^2 + \nu\theta^2 = 2k_i$.

The VG representation provides the null distribution against which the observed integrated statistic $(D_i)$ is evaluated. A large positive $(D_i)$ indicates concordant evidence for upregulation, whereas a large negative $(D_i)$ indicates concordant evidence for downregulation. The corresponding two-sided tail probability measures how unlikely the observed degree of cross-dataset concordance is under the null hypothesis.

**Choice of Parameters**

The derivation above assumes that the signed rank scores contributed by different datasets are independent under the null. In practice, datasets may be correlated because of sample overlap between datasets, batch effects and coupled normalization procedures. Such background correlation increases the variance of the integrated statistics $D_i$, even for features carrying no true biological signal. We therefore used a Brown-style correlation correction[69] in ORBIT to account for dependence between datasets by adjusting the parameters of the VG null distribution.

Importantly, the correlation correction is intended to capture background dependence under the null, rather than biological concordance between datasets. Estimating correlation across all features could therefore lead to over-correction, because genuinely concordant biological signals would contribute to the estimated dependence and are themselves the signals that ORBIT aims to detect.

When individual-level data are available, background correlation $\rho$ can be estimated externally from a permutation null by random permuting the association between molecular measurements and phenotype. When individual-level data are unavailable, we instead estimate correlation $\widehat{\rho_{jj'}}$ from features with no strong evidence of association in either dataset.For each pair of omics layers $(j, j')$, the Pearson correlation, $\rho$, is computed across features observed in both datasets and not called significant in either dataset according to their original per-dataset analysis (adjusted P < 0.05 by default). Residual weak true biological signal may remain in this set and inflate background correlation $\widehat{\rho_{jj'}}$. However, this would produce a wider null distribution and yield conservative rather than inflated significance estimates.

For feature $i$, only the datasets in which the feature is measured contribute to the correlation parameter. Let $A_i$ denote the set of such datasets, we define the pairwise background correlations by their mean

$$\overline{\rho_i} = \frac{2}{k_i(k_i-1)}\sum_{j<j'\in A_i}\widehat{\rho_{jj'}}.$$

This feature-specific form accounts for differences in the datasets contributing to each feature and does not require all pairwise correlations to be equal. If all pairwise correlations are equal, $\overline{\rho_i}$ simply reduces to a common $\rho$, as the per-feature form, remains valid for an arbitrary correlation matrix.

The variance of $D_i$ follows directly from the pairwise correlations, with no distributional assumption beyond the second moment:

$$\mathrm{Var}(D_i) = 2k_i\lambda_i,\ \lambda_i = 1 + (k_i - 1)\bar{\rho}_i.$$

The exact distribution of a sum of correlated Laplace variables is not analytically tractable. We therefore adopt a Brown-style correlation in which correlation is represented by an effective number of independent datasets,

$$\mathrm{k}_{\mathrm{i}}^{\{\mathrm{eff}\}} = k_i/\lambda_i$$

As correlation increases, $k_i^{eff}$ decreases, reflecting the reduced amount of independent information contributed by correlated datasets. This gives the approximate excess kurtosis:

$$\mathrm{Kurt}_{\mathrm{ex}}(D_i) \approx \frac{3}{n_{\mathrm{eff}}} = \frac{3\lambda_i}{k_i},$$

We therefore adjust the VG parameters to match the variance and excess kurtosis of the correlated statistic, giving

$$\nu_i = \lambda_i/k_i = \frac{1+(k_i-1)\bar{\rho}_i}{k_i},\ \text{and}\ \sigma_i = \sqrt{(2k_i\lambda_i)} = \sqrt{2k_i[1 + (k_i - 1)\bar{\rho}_i]}.$$

When $\bar{\rho}_i = 0$, this reduces to the independent-dataset case. In contrast, as $\bar{\rho}_i \to 1$, the datasets become fully correlated and therefore provide only one independent source of information. Because the same score is summed across $k_i$ datasets, $D_i$ approaches a Laplace distribution with scale $k_i$, $\mathrm{Laplace}(0, k_i)$.

**Numerical evaluation of VG tail probabilities.**

Existing implementation of the VG distribution are not designed for extreme tail probabilities required by omics analyses[70]. In particular, the implementation used in [70] returns probabilities below $10^{-10}$ as zero, and provides limited accuracy in extreme tail ($P < 10^{-6}$). ORBIT therefore implements a dedicated log-space evaluator. Because the VG law is symmetric about zero, the two-tailed P value for feature $i$ takes the unified form

$$P_i = P\left(|D_i| \geq |D_{i,\mathrm{obs}}|\right) = 2\int_{|D_{i,\mathrm{obs}}|}^{\infty} f_{D_i}(x; \sigma_i, \nu_i)dx$$

For general $\nu_i$, this integral does not have a simple closed-form solution and is evaluated numerically. Under our moment matching, the exponential tail scale of the density, $\sigma_i\sqrt{\nu_i/2}$, equals the variance-inflation factor $\lambda_i$ defined above, so the branch expressions below are written in terms of $\lambda_i$; we dispatch to one of three branches according to the value of $\nu_i$. (i) Floating-point guard. When $|D_{i,\mathrm{obs}}| < 10^{-10}$, we return 0. (ii) Closed-form special cases. When $\nu_i$ is numerically indistinguishable from 1 or $1/2$, the tail integral is elementary. For $\nu_i = 1$ the law reduces to $\mathrm{VG}(\sigma_i, 1,0,0) = \mathrm{Laplace}(0, \lambda_i)$, giving $\log P_i = -|D_{i,\mathrm{obs}}|/\lambda_i$, and for $\nu_i = 1/2$, integrating the density by parts gives $\log P_i = \log\frac{1}{2} + \log(z+2) - z$ with $z = |D_{i,\mathrm{obs}}|/\lambda_i$. Dispatching to these expressions returns the exact value at negligible cost and avoids quadrature in the two cases where it is unnecessary. (iii) General case. For all other $\nu_i$ we evaluate the tail integral by 32-node Gauss–Laguerre quadrature. The substitution $x = |D_{i,\mathrm{obs}}| + \lambda_i u$ maps the upper tail to the canonical Laguerre form

$$P_i = 2\lambda_i \int_0^{\infty} \left[f_{D_i}\left(|D_{i,\mathrm{obs}}| + \lambda_i u\right)e^{u}\right] e^{-u}du$$

in which the $e^{-u}$ weight is absorbed by the Laguerre nodes, leaving the rescaled integrand $g(u) = f_{D_i}(d_i + \lambda_i u)\, e^{u}$. Because $f_{D_i}$ carries the modified Bessel factor $K_{1/\nu_i - 1/2}$, whose intrinsic decay is $e^{-x/\lambda_i}$, the exponential is cancelled exactly by the rescaling and $g(u)$ retains only a power-law prefactor — precisely the regime for which Gauss–Laguerre quadrature is well suited.
