## Supplementary Figures for "Rank-based integration identifies convergent disease mechanisms across omics"

### Supplemental Figures

**Supplemental Fig. 1. Agreement between the analytical variance-gamma null and Monte Carlo simulation under the continuous-rank model.** Two, three and five omics layers were simulated with normalized ranks drawn as continuous Uniform (0,1) variables and random signs. P_VG is the analytical tail probability as implemented in ORBIT. P_MC was obtained by direct Monte Carlo simulation with $10^{10}$ draws per layer count and is shown with exact 95% Poisson confidence intervals. The dashed line marks a ratio of 1.

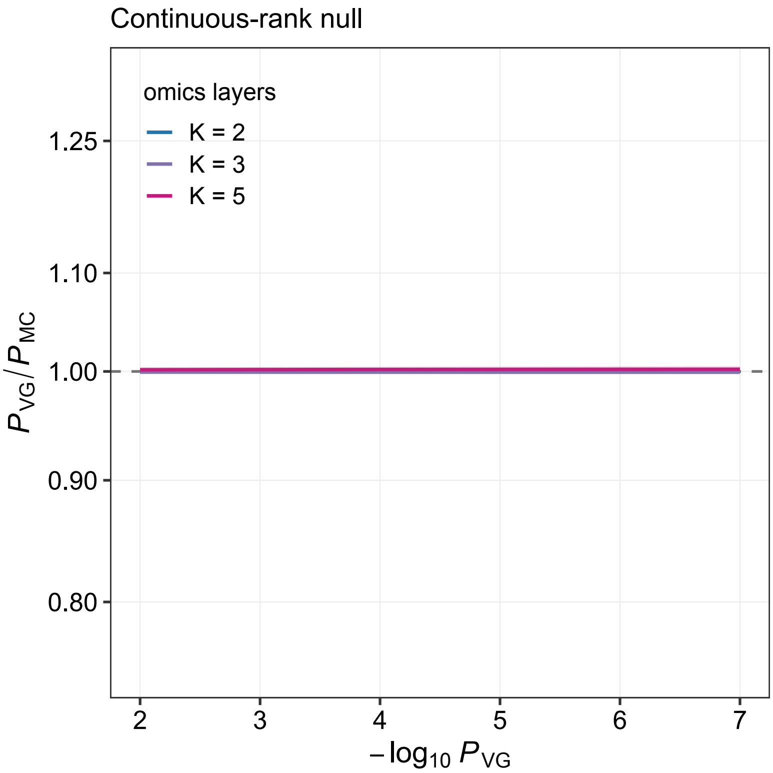

**Supplemental Fig. 2. Conservative bias introduced by the discreteness of the rank transform. (A)** Two omics layers with 5,000, 10,000 and 20,000 features per layer. The rank null is obtained by exact enumeration over all n² rank pairs and four sign combinations, so these curves carry no sampling error. **(B)** 10,000 features per layer, with two, three and five omics layers. The K = 2 curve is again exact; K = 3 and K = 5 were obtained by rank permutation 2 × $10^8$ and are shown with pointwise 95% confidence bands.

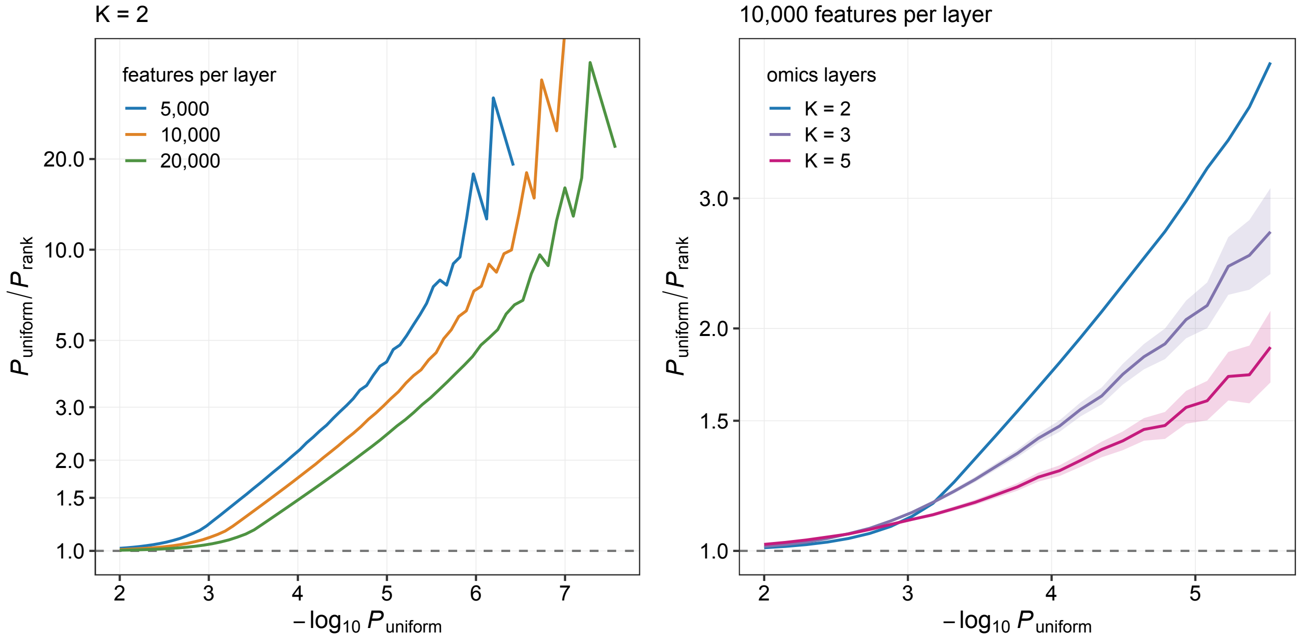

**Supplemental Fig. 3 Power and false-positive rate of RankProd compared with ORBIT under identical simulations. (A-D)** False-positive rate at four proportions of signals (0%, 10%, 20%, 50%) for no signals (solid line), independent signals (dashed line) and opposite signals (dotted line) scenarios. **(A)** Varying between-omics correlation. **(B)** Varying number of omics layers. **(C)** Varying proportions of missing values with median imputation. **(D)** Varying proportions of missing values when missing values were ignored **(E)** Per-feature P values from ORBIT and RankProd on the same simulated data as in **Figure 2C** of opposite-direction signal. **(F)** Power of RankProd and ORBIT to recover concordant signal as a function of the number of omics layers, at five proportions of signals (0%, 5%, 10%, 20%, 50%).

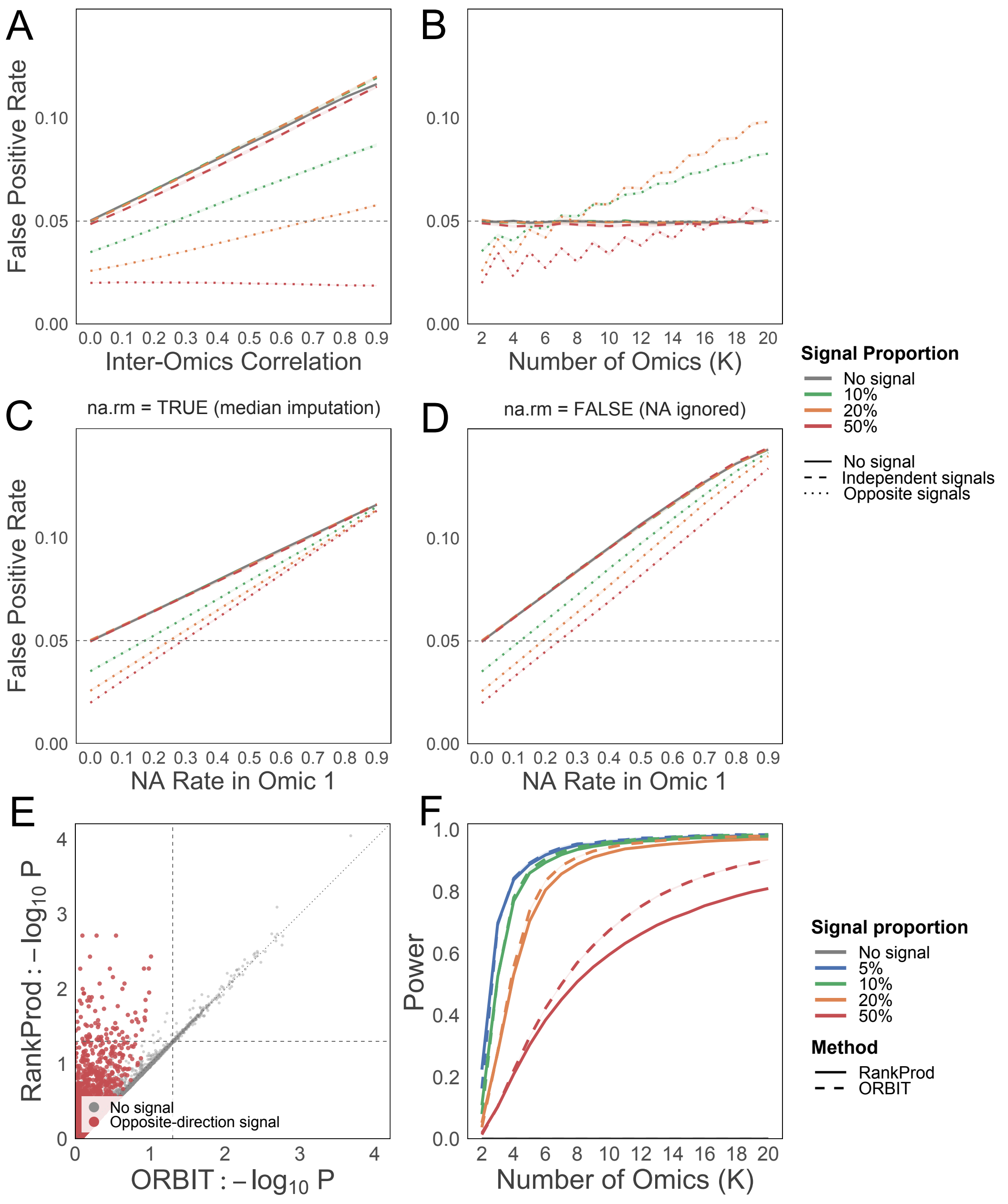

**Supplemental Fig. 4. CKD gene prioritized by Fisher, Stouffer and DPM.** (**A**) The top 10 most significant genes by Fisher's method. **(B)** The top 10 most significant genes by DPM. **(C)** The top 10 most significant genes by Stouffer's method. Asterisks mark genes in **A-C** not among the top 10 under ORBIT.

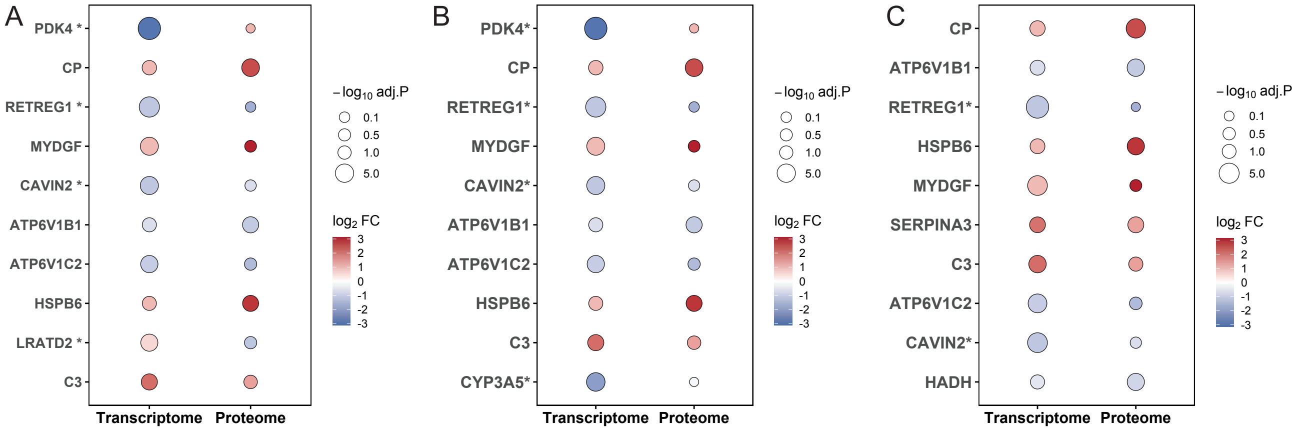

**Supplemental Fig. 5. Comparison of ORBIT and RankProd for features with opposite directions of change across omics layers.** (**A**) Signed significance, sign($\log_2$FC) × $-\log_{10}$P, in the transcriptome versus the proteome for genes measured in both layers. Genes with opposite directions of change that rank within the top 20% by P-value in both layers are highlighted in purple. The dotted line indicates the line of identify (y = x). **(B)** Comparison of $-\log_{10}$P from ORBIT and RankProd for the same genes, with opposite-direction genes highlighted as in (**A**). Dashed lines indicate P = 0.05 and the dotted line indicates the line of identity (y = x).

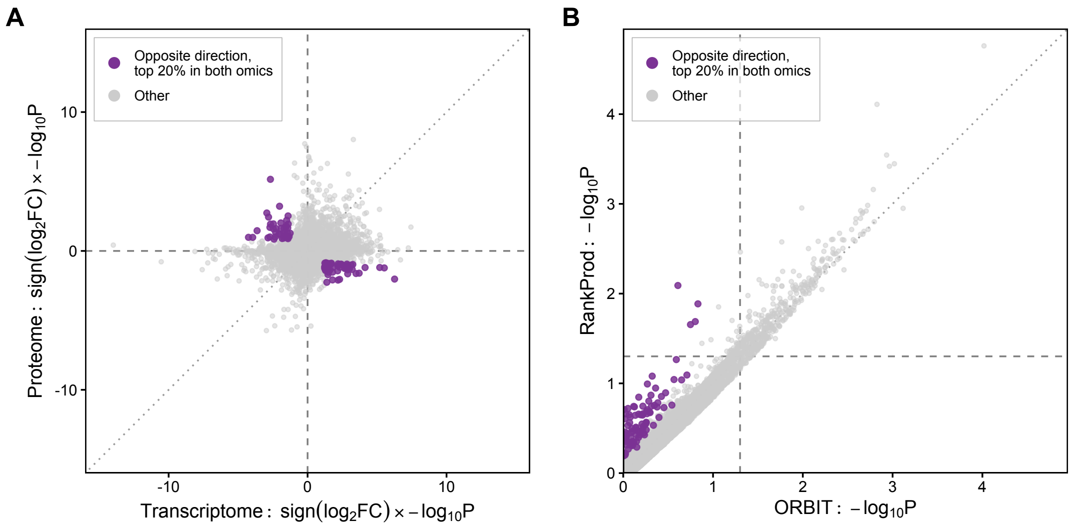

**Supplemental Fig. 6. Comparison of pathway significance under GSEA and over-representation analysis using the integrated ORBIT transcriptome.** Restricted to pathways significant (adjusted P value <0.05) in both methods. Each point is a gene set, positioned by its $-\log_{10}$(adjusted P value) under each approach, colored by direction and sized by the number of member genes recovered in the ORA input set. The top 10 most significant pathways under GSEA are labeled.

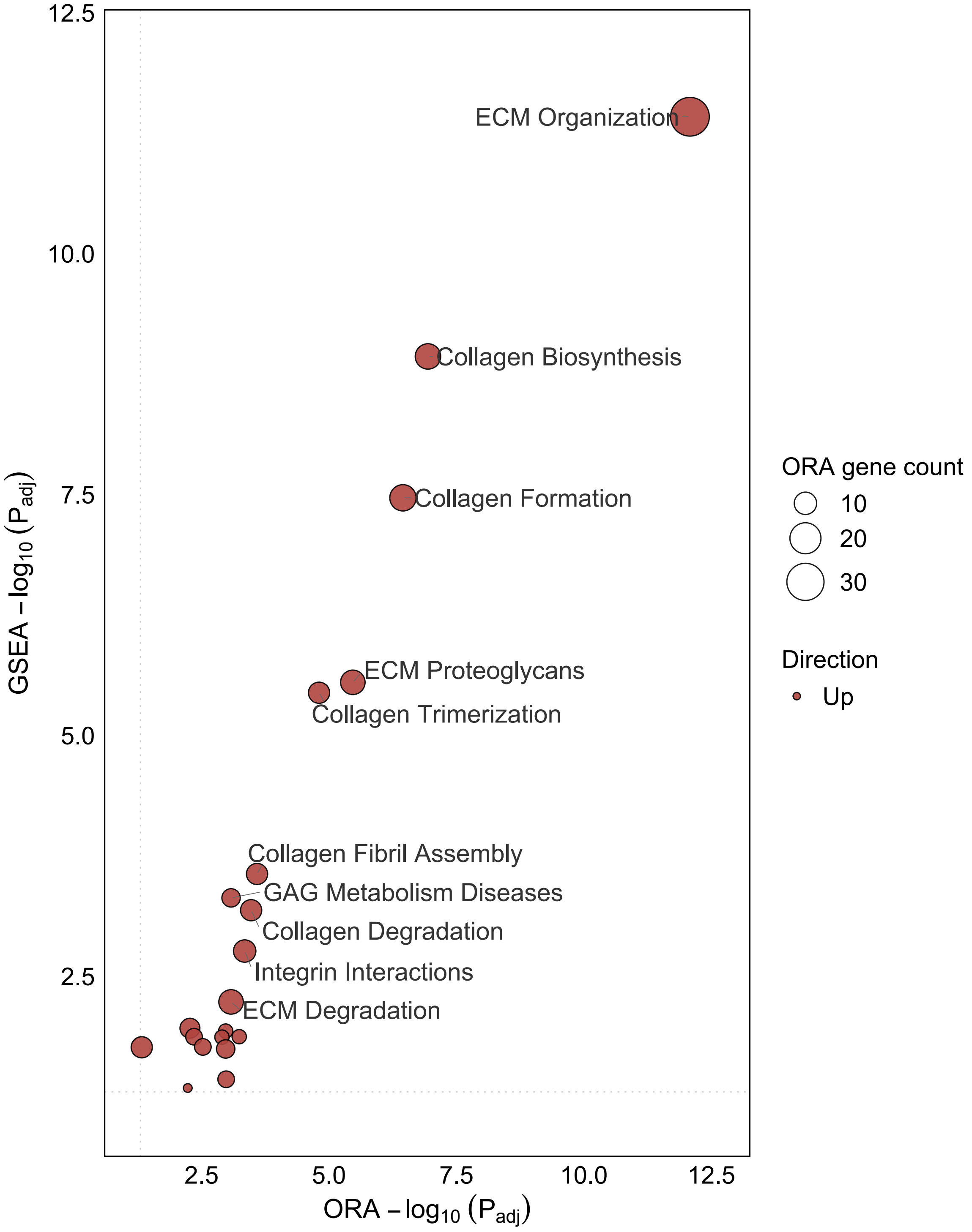

**Supplemental Fig. 7. The top 10 most highly ranked proteins across the four contributing proteomic studies.** Colour indicates $\log_2$(Fold Change) and point size indicates the per-omic $-\log_{10}$(adjusted P value).

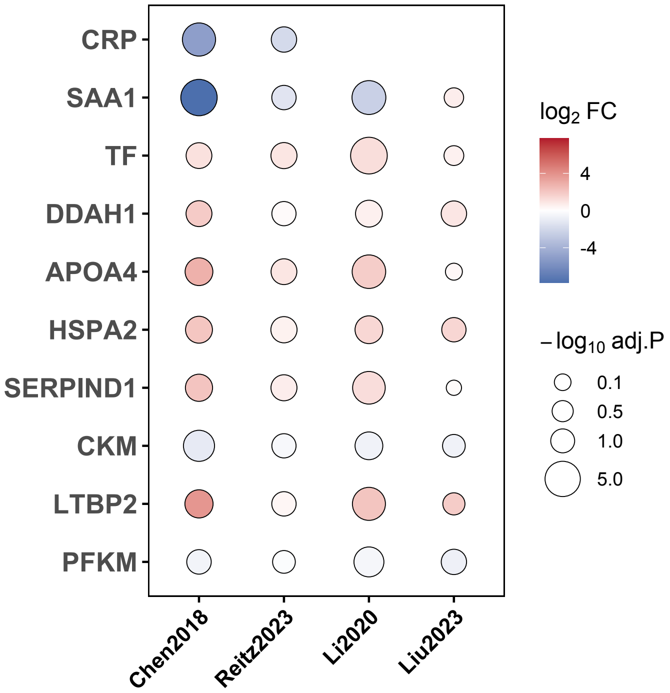

**Supplemental Fig. 8. The false-positive rate control of ORBIT-P in simulation. (A-O)** False-positive rate for no signals. **(A, D, G, J, M)** False-positive rate under varying between-omics correlation. **(B, E, H, K, N)** false -ositive rate under varying omics layers. **(C, F, I, L, O)** false-positive rate under varying proportion of missing values. **(A-C)** Fisher method. **(D-F)** Brown method. **(G-I)** Directional Stouffer method. **(J-L)** Directional P-value Merging (DPM). **(M-O)** ORBIT-P method. **(P)** Power of ORBIT-P (dashed line) and DPM (solid line) to recover concordant signal as a function of the number of omics layers, at five proportions of signals (0%, 5%, 10%, 20%, 50%).

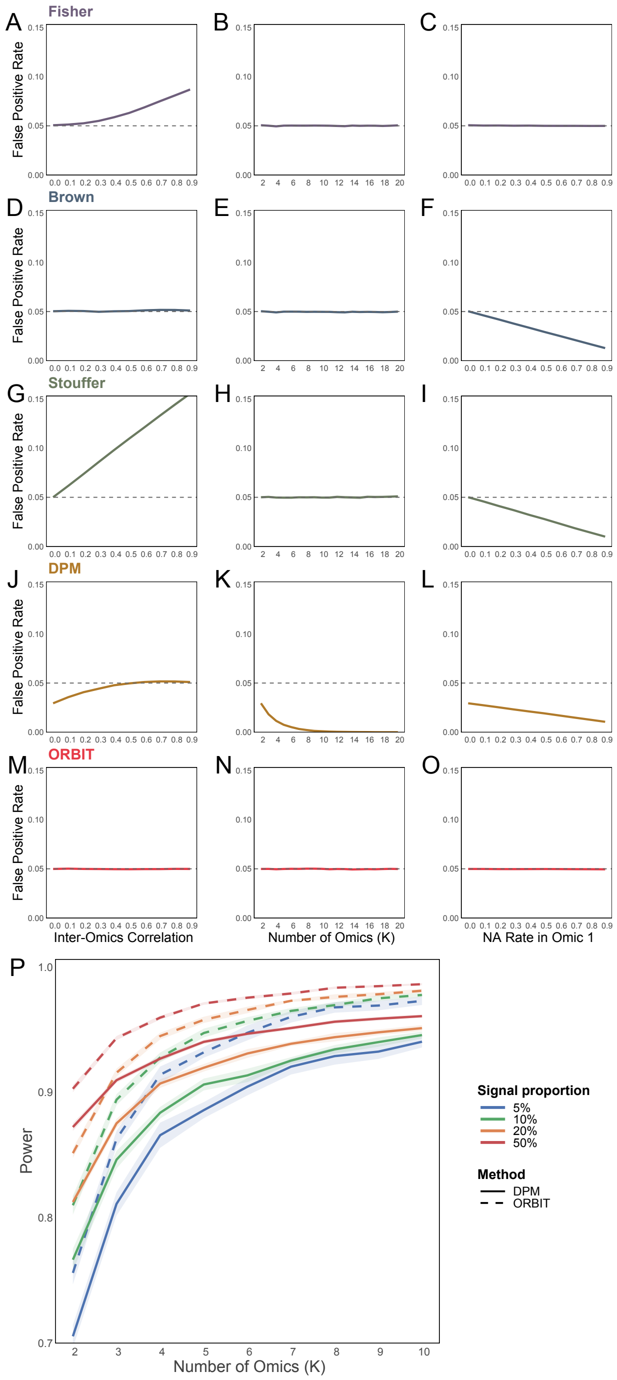

**Supplemental Fig. 9. Monte Carlo validation of the ORBIT and Variance-Gamma Package [1]. (A)** Computed against Monte Carlo tail probability, both on a $-\log_{10}$ scale. The dashed line is the identity. Horizontal bars give the Monte Carlo 95% confidence interval. **(B)** The same comparison expressed as relative deviation from the Monte Carlo estimate. The grey band is the Monte Carlo 95% confidence interval.

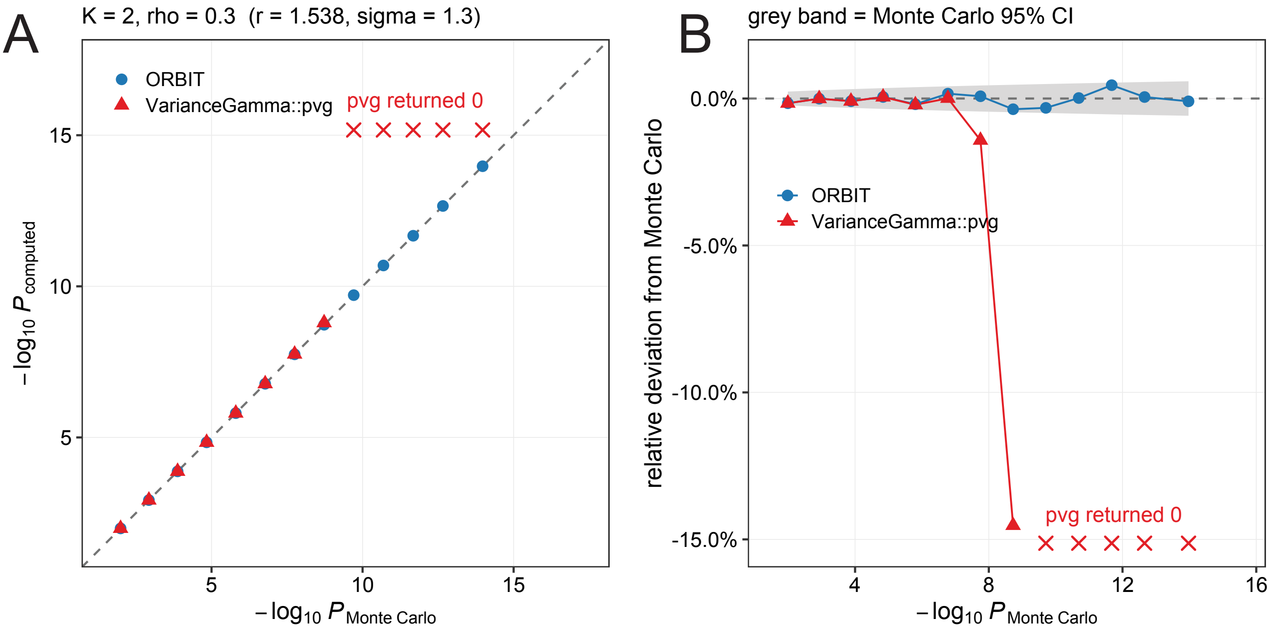


1. Scott, D. and C.Y. Dong, *VarianceGamma: The Variance Gamma Distribution*. 2023.
